# Proof of Concept for High-Dose Cannabidiol Pretreatment to Antagonize Opioid Induced Persistent Apnea

**DOI:** 10.1101/2024.09.13.612358

**Authors:** Beth M. Wiese, Evgeny Bondarenko, Jack L. Feldman

## Abstract

Using a mouse equivalent of FDA-approved cannabidiol (CBD) dosing, we found high dose CBD affects opioid induced persistent apnea (OIPA), the principal cause of opioid related fatalities. CBD pretreatment mitigated respiratory depression from fentanyl in awake mice and significantly delayed OIPA onset in anesthetized mice, effective as the opioid antagonist naloxone. The powerful effect of CBD pretreatment on OIPA suggests a novel therapeutic strategy to reduce fatal opioid overdose incidence.

## Introduction

Opioids activate µ-opioid receptors to produce analgesia, but also depress brainstem circuits that generate breathing movements that can result in a persistent and fatal apnea (1-3). Administration of the opioid antagonist naloxone can restore breathing after onset of opioid-induced persistent apnea (OIPA), but also reverses analgesia, euphoria, and induces precipitated withdrawal in opioid dependent individuals.

Certain cannabinoids, including high dose CBD, have therapeutic benefits in the treatment of substance use disorders, pain, anxiety, and gastrointestinal ailments. Specifically, some cannabinoids can increase breathing frequency in preclinical models of opioid-induced respiratory depression (4, 5), as well as inhibit binding at opioid receptors (6-11), and increase the efficacy of naloxone (6, 12).

Based on FDA approved dosing recommendations for CBD that are well tolerated in humans (9-11, 13-15), we determined a mouse equivalent dose of 250 mg/kg for investigation (16). We predicted CBD pretreatment would attenuate fentanyl-induced respiratory depression in awake mice and delay the onset of OIPA in urethane anesthetized mice.

## Results

### Awake Behaving Mice

We assessed changes in breathing frequency in awake behaving mice resulting from a single fentanyl dose (50 mg/kg i.p.) that followed i.p. pretreatment with saline, vehicle, naloxone (NX; 100 mg/kg), CBD (250 mg/kg), or CBD+NX. There was no impact on breathing frequency following pretreatment injections across groups compared to within-subject baseline (Fig.1A). Fentanyl after pretreatment with saline or vehicle significantly decreased breathing frequency (∼60%; Fig.1B). In contrast, pretreatment with CBD, NX or CBD+NX attenuated this reduction in breathing frequency to 25% or less (Fig.1C). To determine effects of these pretreatments on decreasing the lethality of fentanyl, we examined their effects on breathing frequency during continuous fentanyl infusion in anesthetized mice.

**Figure1.**
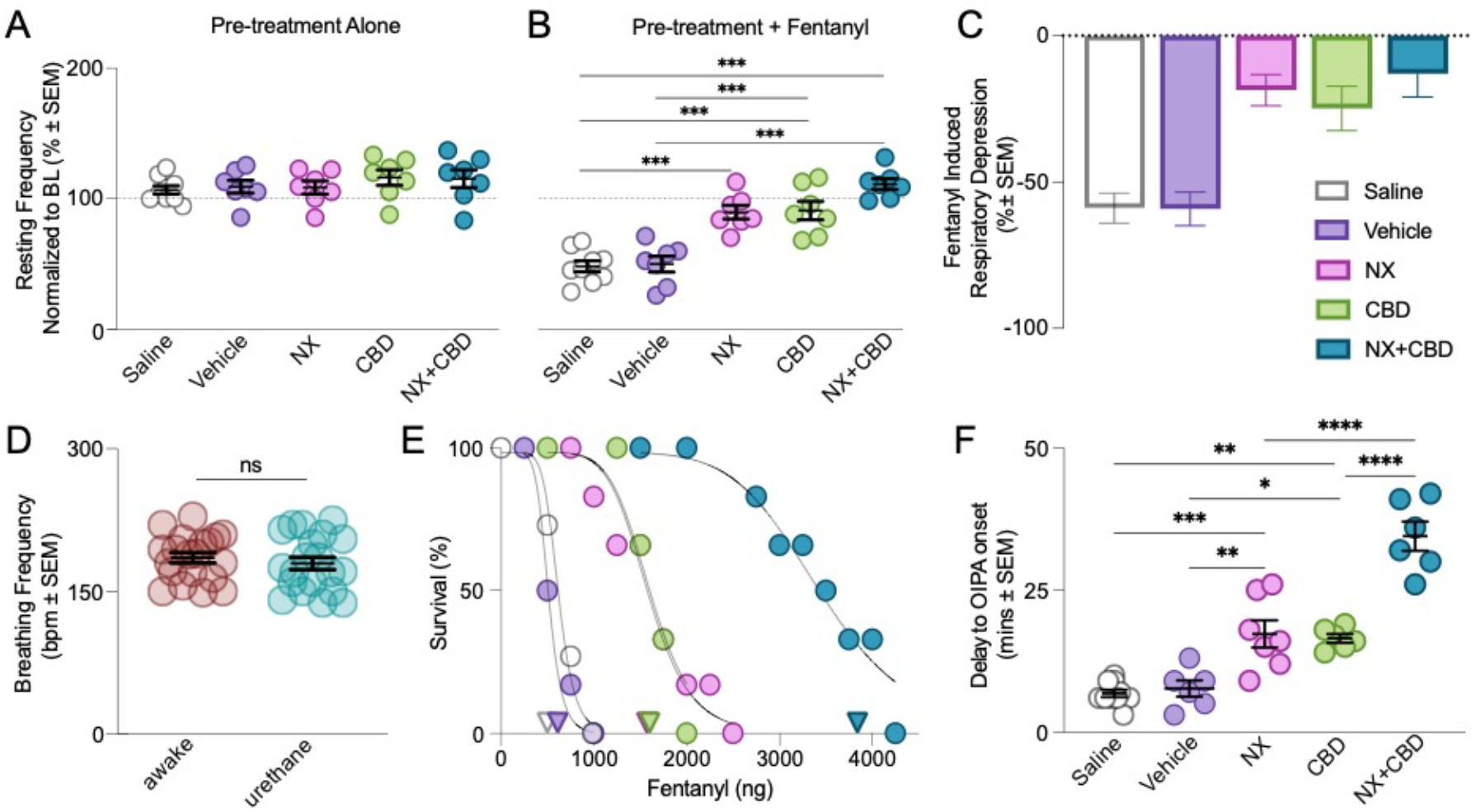
Pretreatments with CBD, NX or both mitigate fentanyl-induced respiratory depression and OIPA in mice. **A** Breathing frequency in awake mice following pretreatment alone was not altered compared to within-subject baseline (BL), (F (32, 64) = 1.36, p = 0.15) or across groups, (F (4, 65) = 0.73, *p* = 0.58). **B**. Significant decreases in breathing frequency are seen in response to fentanyl in awake mice, (F (8, 65) = 13.42, *p* < 0.001), after pretreatment with saline, n=9, *p* < 0.0001, or vehicle, n=7, *p* < 0.001, compared to NX, n=7, *p* = 0.19, CBD, n=7, *p* = 0.27, or CBD + NX, n=7, *p* = 0.38. **C**. Awake mice pretreated with saline or vehicle significantly depressed breathing frequency ∼60%, while NX, CBD, or CBD + NX, only reduced breathing frequency by 25% or less following fentanyl administration. **D**. Average breaths per minute (BPM) in urethane anesthetized mice (n=20) were the same as in awake mice (n=20), *p* = 0.48. **E**. Survival functions (see METHODS) of urethane anesthetized mice during continuous ICV fentanyl infusion were used to determine the LD_50_ of fentanyl after pretreatment (Triangles correspond to LD_50_ values) with saline (480 ng; 95%CI [0.2, 0.84]) and vehicle (625 ng; 95%CI [0.19, 1.3]). CBD (1611 ng; 95%CI [0.99, 6.5]) or NX (1563 ng; 95%CI [1.0, 6.0]) pretreatment significantly increased the LD_50_ of fentanyl, shifting the curve to the right, and pretreatment with CBD+NX further increased the LD_50_ of fentanyl (3842 ng; 95%CI [2.0, 13.0]). **F**. Delay to OIPA onset from start of continuous fentanyl infusion following pretreatment with saline, n=9, or vehicle, n=6, in urethane anesthetized mice induced a persistent apnea within 9 mins (F (4,30) = 44.83, *p* < 0.001); time to OIPA onset was significantly increased after pretreatment with CBD, n=6, *p* < 0.001, or NX, n=7, *p* < 0.001, (∼17 mins each); time to OIPA onset was further increased following pretreatment with CBD + NX, n=6, *p* < 0.001, (∼35 mins).

### Urethane Anesthetized Mice

We chose urethane anesthesia to investigate the lethal fentanyl dose as it preserves awake breathing frequency in mice (Fig.1D), where fentanyl can produce a persistent apnea, i.e., OIPA. Urethane contrasts with many other anesthetics, e.g., isoflurane, that profoundly reduce breathing frequency (17), as well as the potency of opioids to depress breathing. Here, urethane anesthetized mice, following pretreatment with saline, vehicle, NX (100 mg/kg), CBD (250 mg/kg), or CBD+NX, received a continuous ICV infusion of fentanyl (100 nL/min) i.p. We then determined the delay to OIPA onset and the median lethal dose (LD_50_) of fentanyl from survival functions (see METHODS). Following pretreatment with saline, the LD_50_ of fentanyl was 480 ng, and with vehicle was 625 ng (Fig.1E), with onset of OIPA occurring within 9 mins (Fig.1F). There was a significant increase in the LD_50_ of fentanyl following pretreatment with CBD (LD_50_ = 1611 ng) or NX (LD_50_ = 1563 ng) (Fig.1E) and the delay to OIPA onset (∼17 mins; Fig.1F). Furthermore, compared to CBD or NX alone, pretreatment with CBD+NX significantly prolonged the delay to OIPA onset (∼35 mins; Fig.1F) and further increased the LD_50_ of fentanyl (3842 ng) (Fig.1E).

## Discussion

In awake mice, CBD pretreatment was as effective as NX pretreatment at mitigating fentanyl-induced depression of breathing frequency. In anesthetized mice, CBD pretreatment significantly increased the time to onset of OIPA and the LD_50_ of fentanyl, comparable to that of NX. Furthermore, CBD + NX pretreatment was more efficacious than either CBD or NX alone at increasing the LD_50_ and time to OIPA onset. These data suggest high dose CBD may be an effective prophylactic therapy for OIPA by increasing the time before onset, as well as enhance the efficacy of NX, in the event it is still needed, as an additional strategy to save lives.

The proof of principal effect seen here in mice is robust and has a potential immediate impact for public health in reducing opioid related fatalities. The mechanisms high-dose CBD antagonizes respiratory depression and OIPA are currently not known, and with more than 72 known protein targets, including serotonin, opioid and dopamine receptors, as well as TRPV channels (6-11), and an ability to block adenosine transport (9-11, 18, 19), the scope of potential mechanisms at any site that can generate or modulate breathing (1-3) is legion. CBD also modulates opioid binding activity (8, 11, 12), thus, CBD could antagonize the respiratory depressive effects of µ-opioid receptor activation in the breathing central pattern generator, including in the preBötzinger Complex (1-3). Furthermore, the mechanism could be limited to conditions requiring a homeostatic response since CBD had little to no effect on resting breathing frequency. With these possibilities and given that high dose CBD: i) is FDA approved (10); ii) is well tolerated in humans (15) including when given concurrent with intravenous fentanyl administration (9); iii) does not counteract opioid euphoric effects (9), and; iv) is available without a prescription in the US, there are few barriers to translation of therapeutic implementation for people using opioids, licit or illicit. Only clinical trials in human subjects can determine the lowest effective dose, if any, of CBD. This proof of concept using CBD as a prophylactic therapeutic for prevention of fatal OIPA has considerable potential for public health benefit and is well suited for translation into clinical trial.

## Materials and Methods

### Animals and Drugs

Male C57 mice (n=72) bred in-house under 12-hr light/dark cycle had unrestricted access to food and water. Mice were used under protocols approved by the University of California, Los Angeles, Animal Research Committee and UCLA animal care committee (#1994-159-83). Drugs were administered as stock solutions except for CBD that was dissolved in a vehicle of 10% DMSO, 10% Tween-80, and 80% saline.

### Whole-body plethysmography

Breathing frequency recordings in awake and freely moving mice consisted of one hour baseline, 10 mins after administration of CBD (Cayman Chemical; 250 mg/kg), Naloxone (NX; Pfizer, 100 mg/kg), saline (sigma), vehicle (10% DMSO (Sigma), 10% Tween-80 (Sigma), and 80% saline), or CBD + Naloxone (CBD+NX; 250 mg/kg + 100 mg/kg), i.p.; followed by fentanyl (McKesson; 50 mg/kg, i.p.) administration 10 mins after. All drugs were administered with an injection volume of 10 mL/kg. Breathing frequency was determined in all recordings of awake mice as the mode of breaths per minute (BPM) distributed across a histogram of 5 mins intervals (20), then normalized to within-subject baseline as a percentage.

### ICV infusion preparation and OIPA model

Isoflurane anesthetized mice were transitioned to urethane (1000-1500 mg/kg, i.p.), with body temperature maintained by a heating pad and airflow signal continuously recorded. The dorsal side of the skull was exposed, a small hole was drilled (A/P −4.9, M/L 0.0) and the tip of a Hamilton syringe was lowered into the IVth ventricle (D/V - 2.25). CBD (250 mg/kg), naloxone (NX; 100 mg/kg), saline, vehicle, or CBD + NX (CBD+NX; 250 mg/kg + 100 mg/kg), i.p., was given 10 minutes prior to the start of fentanyl infusion. Fentanyl was infused at a rate of 100 nL/min until onset of OIPA. We calculated lethal fentanyl dose and time to OIPA onset, which we defined as a timepoint when all breathing activity stopped, resulting in a fatal persistent apnea, i.e., cessation of breathing for longer than 1 minute.

### Statistical analysis

Airflow signals were acquired, amplified, digitized, stored, and analyzed using LabChart 8 software and PowerLab 8/16 data acquisition system (ADInstruments). Parameters were calculated offline from the airflow signal. Prism (GraphPad Software) and IGOR software (WaveMetrics) were used to perform analysis. ANOVA analyses were used for within-subject and across treatment groups. *Post hoc* tests were used when significant differences between groups were found. Significance was set at p*≤*0.05. Survival functions were created using Prism with a sigmoidal, 4 parameter logistic nonlinear regression model. Each fit had R^2^ > 0.98 for each group.

## Abbreviations

CBD: Cannabidiol
OIPA: Opioid induced persistent apnea
Mins: Minutes
BL: Baseline
BPM: Breaths per minute
i.p.: Intraperitoneal
ICV: Intracerebral vesicular
LD_50_: Lethal dose to 50% of population

## Acknowledgments

This study was supported by National Institutes of Health/National Institute of Drug Abuse (R21DA056740; R21DA054383), Marion Bowen Neurobiology Postdoctoral Grant Program (UCLA), and National Heart, Lung and Blood Institute (R35 HL135779; R35HL135779).

